# Olfactory preferences and chemical differences of fruit scents for *Aedes aegypti* mosquitoes

**DOI:** 10.1101/2025.02.05.636686

**Authors:** Melissa Leon-Noreña, Genevieve M. Tauxe, Jeffrey A. Riffell

## Abstract

Feeding on the sugar of fruits and flowers is vital for mosquitoes and increases their lifespan, reproduction, and flight activity. Olfaction is a key sensory modality in mediating mosquito responses to sugar sources. Previous studies have demonstrated that natural nectar sources from fruits and flowers can vary in attractiveness to mosquitoes, with some sources preferred over others. However, how the attractiveness of different fruits relates to the chemical composition of their scent and the responses they evoke from the mosquito’s peripheral olfactory system, is still not understood. In this study, we use closely related fruit species and their varieties to examine how changes in scent chemistry can influence the fruit’s attractiveness to *Aedes aegypti* mosquitoes and examine how the mosquito’s olfactory responses (via electroantennogram recordings, or EAG) correlate with those differences. Our results show that mosquitoes are attracted to the scents of certain fruits (*Mangifera indica, Prunus perspica, Psidium guajava, Musa acuminata*), whereas others (*Pyrus communis, Citrus limon*) elicited responses not significantly different from the negative control. Chemical analyses of the scents showed that attractive fruits have distinct chemical profiles, and amongst closely related fruits, minor changes in the relative proportions of scent compounds can modify the attractiveness. These minor differences in the fruit scent were not reflected in the EAG responses, which showed similar responses to scents from different fruit species and closely related varieties. Experimentally altering the chemical proportion of a single compound in attractive scents caused a significant decrease in attraction to levels similar to the less attractive cultivars. Our results demonstrate that mosquitoes are sensitive to compound proportions in attractive odors, which have implications for the olfactory processing of complex odor sources, like those from plants or blood hosts.

**Summary Statement:** *Aedes aegypti* mosquitoes show specific and selective fruit scent preferences related to differences in the proportion of compounds in the scent.

## Introduction

Plant-sugar feeding is a critical component for adult mosquitoes, with both males and females utilizing sources of sugar from floral, fruit, and extrafloral sources throughout their lives (Bradshaw et al., 2018; Brantjes and Leemans, 1976; Foster, 1995; Gu et al., 2011; Jhumur et al., 2007; Lahondère et al., 2020; Okech et al., 2003; Peach and Gries, 2016; Yuval, 1992). Plant sugar is an essential part of the mosquito diet and the only source of food for males. Although females can use nutrient sources from blood meals, sugar from plants is still critical for other metabolic and behavioral processes, such as flight and oviposition (Foster, 1995; Yuval, 1992). Previous studies have shown that mosquitoes exhibit behavioral preferences to certain flowers and fruits, with some preferred over others (Manda et al., 2007; Müller et al., 2011; Nikbakhtzadeh et al., 2014; Yu et al., 2017). For example, in western Kenya, *Anopheles gambiae* mosquitoes exhibited distinct preferences to flowering plants, ranging from strong attraction to repellency or neutral behaviors (Manda et al., 2007; Müller et al., 2010). Similarly, in field experiments, *Aedes albopictus* mosquitoes were selectively attracted to certain, diverse flower species and rotting fruits (Müller et al., 2011), and in laboratory experiments with *Aedes aegypti* mosquitoes have shown clear preferences for specific flowering plants that also serve to increase their longevity (Chen and Kearney, 2015). However, the mechanisms by which mosquitoes discriminate between sources of plant sugar are not clear.

Olfaction is a key sensory modality mediating the adult mosquitoes’ ability to locate sources of food, including blood (De Obaldia et al., 2022; Zwiebel and Takken, 2004) and sugar (Nikbakhtzadeh et al., 2014; Vargo and Foster, 1982; Von Oppen et al., 2015). Although ongoing work is shedding light on the relationship between human scent differences and relative attractiveness in mosquitoes (De Obaldia et al., 2022; Giraldo et al., 2023), few studies have examined comparative differences in the scent chemistry of plant sugar sources and identified electrophysiologically active compounds in the scent (Barredo and DeGennaro, 2020; Jhumur et al., 2007; Lahondère et al., 2020; Nikbakhtzadeh et al., 2014; Nyasembe et al., 2018; Upshur et al., 2023). Flowering plants and fruits that are attractive to *An. gambiae* mosquitoes have been shown to emit electrophysiologically active compounds, including monoterpenes, sesquiterpenes, and aliphatic compounds (Meza et al., 2020; Nyasembe et al., 2018). *Ae. aegypti* are selective in their antennal responses to monoterpenes, sesquiterpenes, esters, and aromatics (Lahondère et al., 2020; Nyasembe et al., 2018). However, the fruits and flowers that were less attractive, or neutral in their attractiveness, emitted many of the same compounds. It remains unclear which features of the scent – such as composition or intensity – may be driving these behavioral differences.

As odors are transported from the sources, their concentrations vary in space and time due to turbulent mixing by the wind (Riffell et al., 2008). Insects, including mosquitoes, can recognize behaviorally relevant odor sources despite these fluctuations in intensity (Dekker and Cardé, 2011; Murlis et al., 1992). For many insects, the proportion of certain key compounds in the scent is critical for recognizing attractive odor sources as the plume fluctuates in concentration, providing a chemical fingerprint for searching insects (Lahondère et al., 2020; Martin and Hildebrand, 2010). Examples of this phenomenon comes from diverse insect species, including the sex pheromone system in Lepidoptera, where female moths emit a sex pheromone mixture of two to three key compounds at specific concentrations, the proportions of which are critical for the recognition by searching males (Martin et al., 2013; Roelofs and Cardé, 1977). Similar results have been shown in mosquitoes, where a floral species (*Platanthera obtusata*) attractive to *Aedes spp.* mosquitoes emitted scents dominant in aliphatic aldehyde compounds (e.g., nonanal, and octanal) and low levels of monoterpenes (e.g., lilac aldehyde), whereas a sister species (*P. stricta*), pollinated by bees or moths, emits a fragrance similar to *P. obtusata* but dominated by lilac aldehyde, which is repellent to mosquitoes (Lahondère et al., 2020).

Differences in the proportions of various odorants in complex scents could explain the variation in mosquito attraction to sources of plant sugar. The proportion of compounds in the scent of fruits differs widely, including those from closely related genera and even cultivars of the same species (Bouzayen et al., 2009; Du and Ramirez, 2022; Li et al., 2021). For example, the date palm – a favored fruit in mosquito lures – has different fruit cultivars that overlap in their scent composition, with some having a higher concentration of repellent terpenoid compounds, such as citronellol, whereas others have a higher concentration of attractive compounds, such as aliphatic aldehydes (Guido et al., 2011; Khalil et al., 2017). However, a systematic examination of the scents between closely related species and their relative attractiveness to mosquitoes has yet to uncover the relative importance of different scent features (composition, concentration, or proportions) in attracting mosquitoes.

In this study, we take advantage of closely related fruits and their cultivars to examine the relation between the scent composition, antennal olfactory responses, and the scent attractiveness to *Aedes aegypti* mosquitoes. Whole ripe and overripe fruits are an attractive and important sugar source for mosquitoes, and mosquitoes have been shown to pierce the fruit peels to access the sugar and plant nutrients (Müller et al., 2011). Testing different fruit species—including those used in mosquito lures—and those of different cultivars allowed us to examine how scents overlapping in composition can evoke different levels of behavioral attraction. We present findings from 1) behavioral tests of different fruits and fruit varieties or cultivars, 2) analyses of fruit scent volatile compounds and emissions and their sources, 3) electrophysiological responses of the mosquito antennae to the fruit scents, and 4) behavioral experiments showing how changes to a compound proportion in the fruit scent alter mosquito attraction. Using this integrative approach, we demonstrate that, for *Ae. aegypti*, attraction to fruit scents depends upon the chemical composition and the proportions of the scent, which has important implications for the olfactory processing of complex odors in *Ae. aegypti* mosquitoes, and future development of attractive lures.

## Methods

### Mosquito rearing

*Ae. aegypti* mosquitoes used for behavior experiments were provided by BEI Resources (Manassas, VA, USA) and reared at the University of Washington (Seattle, WA, USA). In preliminary experiments, different *Ae. aegypti* lines (Rockefeller, Liverpool, Costa Rica, and Puerto Rico, all from BEI Resources) were tested in their response to scent from mangoes (*Mangifera indica* ‘Tommy Atkins’), and all showed qualitatively similar levels of attraction with approximately 55% to 85% of the mosquitoes attracted to the scent. Although the tested mosquitoes have remained in various insectaries for many generations, the responses to mango and other attractive fruits may suggest the fruit scents evoke an innate behavioral response. For the remainder of the experiments, we used the Rockefeller line which showed consistent and robust responses to the fruit scent. Mosquitoes were maintained in an ACL2 insectary, per University of Washington Biological Use Authorization (BUA# 0530-003), at 27°C, 70-80% RH, and a photoperiod cycle of 12h light/12h dark. Eggs were hatched in plastic trays and deoxygenated with deionized water. Groups of 200 larvae were placed in covered trays containing tap water and fed with fish food (Hikari 129 Tropic 382 First Bites - Petco, San Diego, CA, USA). Pupae were grouped based on similar age and isolated in 16 oz containers (Mosquito Breeder Jar, BioQuip® Products, 131 Rancho Dominguez, CA, USA) and allowed to emerge. Experiments were conducted using adult mated mosquitoes 6-7 days old and fed 10% sucrose until 24 hours before behavior experiments. Female mosquitoes were not blood-fed.

### Fruit selection

Fruits were selected due to prior work on attracting mosquitoes and their use in mosquito toxic sugar baits, their presence in tropical and subtropical regions with endemic mosquito populations, and their availability to the study. We used intact fruits rather than fruit juices, concentrates, or syrups that do not reflect the natural scent emissions experienced by the mosquitoes (Joseph, 1970). Several of these fruits are similar in species or variety to those used in traps, and their corresponding studies and sources can be found in Table S1. The following fruit species and associated varieties or cultivars were tested: 1) mangoes: *M. indica* ‘Ataulfo,’ ‘Tommy Atkins,’ and ‘Keitt,’; 2) guavas: *Psidium guajava* ‘Pink’ and ‘White’; 3) plums: *Prunus salicinia* ‘Santa Rosa,’ and ‘Burgundy,’; 4) peaches: *P. persica* ‘White Lady,’ and ‘Monroe’; 5) nectarines: *P. persica variety (var) nucipersica* ‘Fantasia,’ and ‘Snow Queen’; 6) bananas and plantains: *Musa acuminata* ‘Cavendish,’ and *M. x paradisiaca*; 7) pears: *Pyrus communis* ‘Williams,’ and *‘*Korean*’*; 8) date palms: *Phoenix dactylifera* ‘Barhi’ and ‘Medjool’; 9) tomatoes: *Lycopersicon esculentum;* and 10) lemons: *Citrus limon*. Fruits selected for each experiment were ripe and were inspected to have no signs of mold, bruises, or damaged skin.

### Two-choice behavior assay

To measure the preference of female and male mosquitoes towards fruit odors, a two-choice behavior assay was created consisting of a cage (Bugdorm, 60cm x 60cm x 60cm) with two smalle traps (35 cm long) placed inside (Fig 1A). Both traps contain an opaque chamber where the whole fruit or control odor source (10% sugar cotton ball) was placed. This chamber was connected by a polytetrafluoroethylene tube to a funnel trap, allowing scent from the fruit or control to passively enter from the first chamber to the trap. The traps were set on opposite sides of the cage, and the placement of traps was randomized between replicates. Mosquitoes were starved of sugar, with water provided via soaked cotton balls 24 h before testing. Once released into the cages, mosquitoes were allowed to choose between experimental and control traps for 48 hours. For each experimental replicate, approximately 75 mosquitoes were placed in each cage. The relative humidity (RH) from each trap was measured using a Sensirion 403-SEK-SENSORBRIDGE (Mouser Electronics, USA) to ensure that the differences in RH between the experimental and control traps did not correlate with an increase in mosquito attraction (r =0.14, p =0.62). After each replicate, trap parts were disassembled and cleaned with 70% alcohol. Fruits were washed with an odorless soap (Tergazyme, Alconox Inc., USA) and allowed to air dry before each experiment. After 48 hours, the total number of mosquitoes in each trap was counted and used in the statistical analyses. Mosquitoes that did not choose between traps were not included. For lemons and tomatoes, 3 replicate trials were performed, while experiments with all other fruits were replicated 6-9 times (Fig1B). Negative control trials (no scent; both traps only contained cotton balls with sugar) were run in parallel with each replicate.

**Figure 1.**
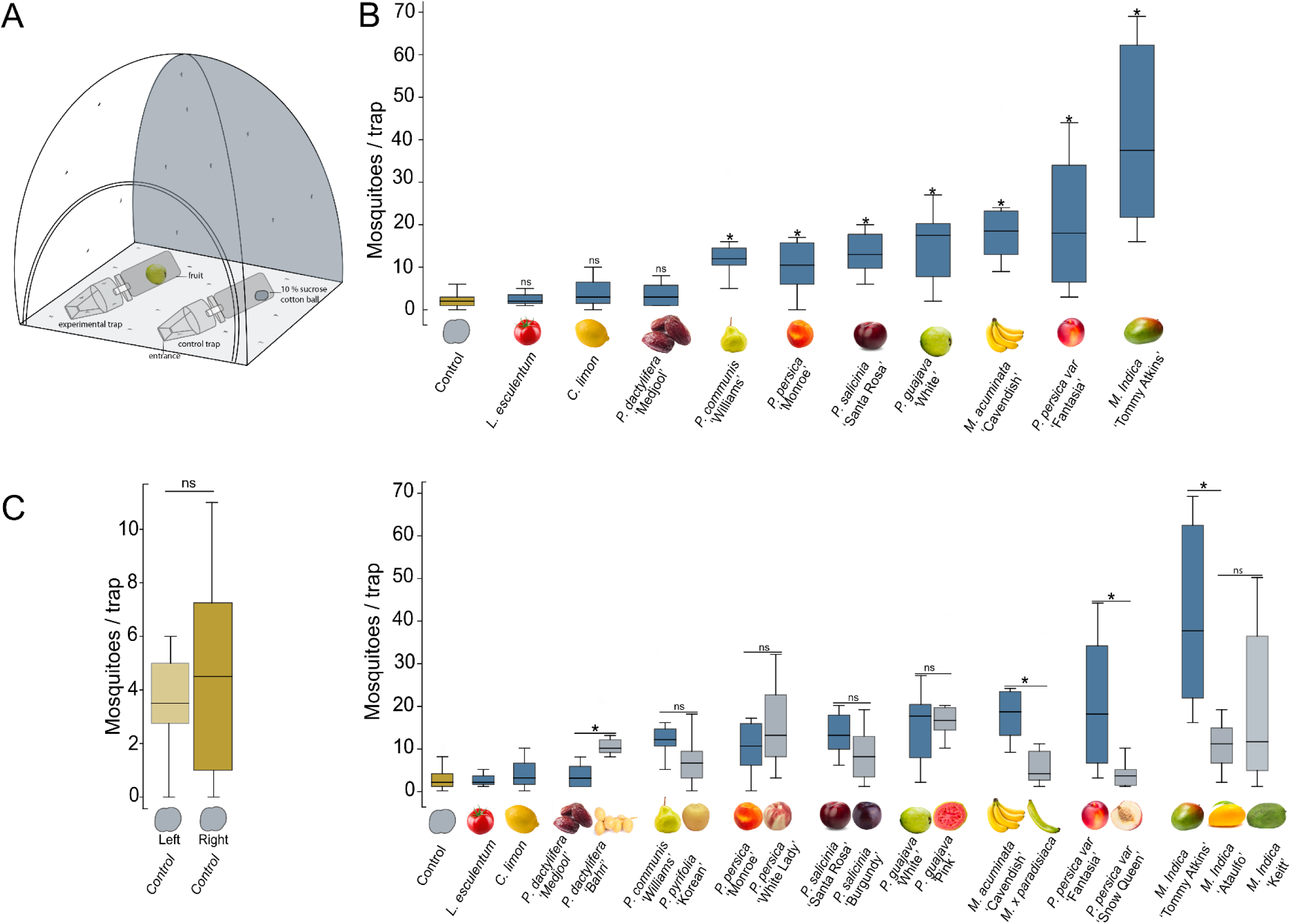
Behavioral preferences of *Ae. aegypti* to fruit scents. **(A)** Schematic of a two-choice behavior assay with control (10% sucrose cotton ball) and experimental trap. **(B)** The total number of mosquitoes inside the experimental fruit trap when provided with a choice between the fruit and control trap. *Top:* Differences in behavioral attraction to different fruit species. The control (n = 9) represents the total number of mosquitoes counted inside the control trap across all fruits in the corresponding graph. *Bottom:* Results testing the differences between closely related species and varieties and cultivars show significant variation in their attractiveness. The control (n = 17) represents the total number of mosquitoes counted inside the control trap across all fruits in the corresponding graph. Bars are the mean ± SE. Asterisks denote a significant difference between each fruit species, variety, or cultivar (Mann-Whitney U-test: *p*<0.05). **(C)** The number of mosquitoes attracted for the negative control trap ran in parallel for each experimental trap with a 10% sucrose cotton ball tested against a 10 % sucrose cotton ball. No significant difference was measured between the two traps (Mann-Whitney U-test: *p*<0.05).

### Fruit headspace collection

Headspace collections were performed to identify volatile organic compounds (VOCs) emitted from each fruit. Each intact fruit was washed with odorless soap (Tergazyme) and air-dried before scent collection. The intact fruit was then weighed and placed inside a nylon bag (Reynolds, USA for 24 hours for volatile collection). Two PTFE tubes (¼” ID X 5/16” OD, fluorostore) were inserted into the bag: one provided air through a charcoal-filtered Pasteur pipette into the bag (1 L/min), and the other vacuumed air from the bag (1 L/min) into a Borosilicate Pasteur pipette containing 100 mg of Porapak powder Q 80-100 (Waters Corporation, USA) and deactivated glass wool (Thermo Fisher Scientific, USA).

VOCs were eluted from each sorbent cartridge in 600 ul of hexane (99%, Sigma-Aldrich, Inc., MO, USA) and stored in 2 ml amber vials at −80° C until analysis. Although Porapak Q does not efficiently capture small molecules like CO_2_ or volatile fatty acids, it does capture diverse volatile compounds greater than 100 Da including monoterpenes, aromatic, aliphatic aldehydes, alkanes, esters, and short-chain alcohols. A series of preliminary experiments were conducted to maximize the capture of diverse compounds, where fruit headspace collections were conducted for different lengths of time (4, 8, 12, and 24 h) and using different amounts of Porapak Q (30, 50, 100, and 200 mg). The 24-hour period, using 100 mg of Porapak Q, enabled us to capture the greatest diversity of VOCs across different fruit species.

For the peel and mesocarp scent collection, an equal weight of each fruit part was placed inside a nylon bag and sampled for 24 hours. Briefly, the fruit peel was manually removed from the mesocarp using a peeler (Oxo Good Grips Y Vegetable Peeler, Oxo Corp., New York, New York) and weighed. The mesocarp was then sectioned to be the same mass as the peel before headspace sampling.

For every series of sample collections, a negative control (empty bag) was run in parallel. In addition, contaminants from the solvent, the sample matrix, and the GC column, were also identified and removed from the datasets.

### Fruit scent chemical identification and quantification

Analyses were conducted in Agilent 7890A GC and 5975C Network Mass Selective Detector (Agilent Technologies, Santa Clara, CA, USA). A DB-5MS GC column (J&W Scientific, Folsom, CA, USA; 30 m, 0.25 mm, 0.25 μm) was used with helium as a carrier gas at a constant 1 cc/min flow. Automated injections of 3 ul for each sample were inserted into the MS using a GS 7693 autosampler (Agilent Technologies, Santa Clara, CA, USA) in spitless mode (220°C) with the oven temperature set at 45℃ held for 6 mins, followed by a heating gradient of 45℃ to 220℃ at 10℃/min, which was then held isothermally for 6 min. Chromatogram peaks were then manually integrated using the ChemStation software (Agilent Technologies, Santa Clara, CA, USA), filtered, and tentatively identified by the online NIST library with confirmation matches >70%. Putative identifications were verified by calculated Kovats Retention indices and comparison to synthetic standards. The concentrations of compounds of interest were determined by comparison to standard curves of synthetic standards measured from 0.5 ng/µl to 1 µg/µl. Total scent emission rates (ng/h) were determined from the quantified scent compounds and normalized to the mass of each fruit. Relationships among the fruit samples’ odor composition were plotted and analyzed using a Non-metric Multidimensional Scaling (NMDS) analysis.

### Electroantennogram experiments

Electroantennograms (EAGs) were performed using similar procedures to Lahondere et al., 2019. Antennae were prepared by dissecting the *Ae. aegypti* heads from the insect and removing the distal tip with tenotomy scissors. The head was placed on the reference electrode with the antennae tips placed on the tip of the Syntech EAG recording probe using Spectra 360 electrode gel-filled (Parker Labs, Fairfield, NJ, USA) so that the electrodes could measure electrical activity moving across the antennae. The EAG electrodes and antennae were placed before a continuous air stream (1000 mL/min flow; Gilmont flowmeter, Gilmont Industries/Barnant Company, Barrington, IL, USA) at 25° C room temperature. The electrodes were connected to a Syntech headstage, connected to an IDAC-4 (Ockenfels Syntech GmbH), allowing 60 Hz noise reduction and filtering. Antennal deflections were counted as responses for a fruit scent if they were 1.5 standard deviations above the noise floor of the antennal activity and occurred within a 0.5-second window of the odor release. The threshold was hand-individually calibrated based on differing levels of signal and noise in each preparation. For each odor stimulus, 8-26 mosquitoes were tested using 5-7 day-old female *Ae. aegypti* mosquitoes. As in previous studies (Riffell et al., 2013; Lahondere et al., 2019), Pasteur pipettes containing the scent extracts were prepared by aliquoting 50 µL onto a small piece of filter paper (Whatman Inc., Clifton, NJ, USA). The hexane solvent was allowed to dry for 7 minutes before the filter paper was inserted in a Pasteur pipette to deliver the fruit scent. Each fruit scent and control stimulus (hexane solvent control, and positive control stimulus 3-methyl-1-butanol [hereafter, isopentanol], diluted at 1% v/v in hexane) was presented randomly. EAG response amplitudes were quantified offline using the Autospike software and normalized to the positive control stimulus.

### Statistical analysis

Statistical analyses were conducted using the Matlab software, v2020b (Mathworks, Natick, MA, USA). The response variable for the behavioral preference assays was the number of mosquitoes in each trap. Non-parametric Mann-Whitney U-tests were deemed suitable, given the lack of normality for within-genera and within-species comparisons. A significant criterion of 0.05 was used for all statistical testing, except those involving multiple comparisons where the criterion was adjusted. The preference assays’ dataset was compared using the total number of mosquitoes per trap using a general linear model (v2020b, Mathworks, Natick, MA, USA). A Kruskal-Wallis test was used to statistically test the relationship between the compounds identified in the fruit scents and the attractiveness of the fruits and to compare the scent emissions.

Non-metric multidimensional scaling analyses (NMDS) were performed to analyze variations in scent composition among fruit varieties and species. For these multivariate analyses, we first coded all identified compounds as either present (1) or absent (0) to examine the dissimilarity between fruits and then constructed a matrix of Bray–Curtis dissimilarities calculated on the relative proportions of the scent compounds. An analysis of similarity (ANOSIM) was performed on the proportion data used in NMDS. ANOSIM is a non-parametric permutation analysis used to assess the similarity between multiple groups regarding the compounds within the scent. To evaluate the clustering in the NMDS, an iterative k-means clustering was performed on the proportional dataset. The number of clustering centroids was determined using the elbow method via computing the distortions under different cluster numbers, where the best cluster number corresponded to 90% of the variance explained (defined as the ratio of the between-group variance to total variance).

## Results

### Behavioral response to fruit scents

As the first step in examining differences across fruit scents, we examined mosquito responses to the negative (no scent) control, run in parallel for each treatment and replicate trial (Fig 1). Across all two-choice behavioral trials, there was no significant difference in the number of mosquitoes attracted to the control trap, or cages containing two control traps (General linear model: t(1,92) = −0.19, *p* = 0.842). On average, 2.4 mosquitoes per trial (±0.25 SE) were attracted to the control trap. By contrast, across all the fruits tested, there were significantly greater numbers of mosquitoes in the baited traps containing the fruits than in the control traps (General linear model: t(1,184) = 6.30, p < 0.001), with 14.0 mosquitoes per trap (±1.2 SE).

There was significant variation in the attractiveness between fruit scents (Fig. 1B). At the species level, mango (*M. indica*) was the most attractive, with a mean of 24.0 mosquitoes per trap (±5.1 SE). Guava (*Psidium guajava*), banana (*Musa acuminata* ‘Cavendish,’), and stone fruits (*Prunus persica*) elicited similar levels of attraction, with approximately 15 mosquitoes per trap (±1.8 SE). By contrast, pear (*Pyrus* spp.) and date palm (*P. dactylifera*) scents, with 10.9 (±2.2 SE) and 8.7 (±1.6 SE) mosquitoes per trap, respectively, were less attractive. Tomato (*L. esculentum*) and lemon (*C. limon*) were the least attractive (2.6 [±1.2 SE] and 4.3 [±2.9 SE] mosquitoes per trap, respectively), and not significantly different from the negative controls (Mann-Whitney U-test: *p* = 0.66).

To examine how scents from closely related fruits may differ in their attractiveness, we tested the cultivars, varieties, and closely related species of fruits (Fig 1B). We again observed significantly different results at the species level (General linear model, species fixed effect: t(3,97) = −9.57, *p* < 0.0001). We also observed significant differences when we examined attraction at the level of varieties and cultivars (General linear model, variety fixed effect: t(3,97) = 15.48, *p* < 0.0001). For example, the scent of the red mango (*M. indica*, ‘Tommy Atkins’) was significantly more attractive than that of the yellow mango (*M. indica* ‘Ataulfo’) (Mann-Whitney U-test: *p* = 0.004). There was a similar effect in the nectarines (*Prunus persica var nucipersica*), with the white nectarine (‘Snow Queen’) attracting four-fold more mosquitoes than the yellow nectarine (‘Fantasia’) (Mann-Whitney U-test: *p* = 0.03). Other pairs of cultivars in the *Prunus* group were not significantly different from one another (Mann-Whitney U-tests: *p* > 0.25)(Fig 1C). There was also a significant difference in the attractiveness of the banana (*Musa acuminata* ‘Cavendish’) and plantain (*M. **×** paradisiaca)* scents (Mann-Whitney U-test: *p* = 0.04), with the banana attracting almost twice as many mosquitoes as the plantain (21.9 and 9.8 mosquitoes/trap for *M. acuminata* and *M. **×** paradisiaca*, respectively), and between the date palm cultivars (*P. dactylifera* ‘Bahri’ and ‘Medjool’; Mann-Whitney U-test: *p* = 0.01). There was no difference in the numbers of attracted mosquitoes for the guava cultivars (*P. guajava* ‘Pink’ vs ‘White’) and between pear species (*P. pyrifolia* vs. *P. communis*)(Mann-Whitney U-tests: *p* = 0.32 and *p* = 0.05 for *P. guajava* and *P. communis*, respectively).

In these experiments, we simultaneously tested male and female mosquitoes. We found there were no significant differences between these two sexes in their relative attraction to the fruit scents (Mann-Whitney U-test: *p* = 0.22), although slightly higher numbers of female mosquitoes were attracted to the fruit scent traps overall (ratio of 1:1.2 male-to-female). Additional control experiments compared male- and female-only trials to those with both sexes and showed no significant difference between experiment types in their numbers of attracted mosquitoes to the fruit traps (Mann-Whitney U-test comparing male-only vs. both sexes: *p* = 0.54; Mann-Whitney U-test comparing female-only vs. both sexes: *p* = 0.78).

### Chemical analysis of fruit scents

VOCs were identified and average emission of fruit scent was determined for each of the nineteen fruit varieties used in behavioral tests. Across all samples, we identified 150 compounds, including 30 terpenoids, 20 aromatics, 2 sulfur, 2 furans, and 96 aliphatic compounds, were identified across all sampled species, varieties, and cultivars (Table S2). There were significant differences in the emission rates and total number of scent compounds among the fruit samples (Kruskal-Wallis test: χ16,74 > 53.84, p < 0.001), but there was no significant correlation between these factors and the fruits’ attractiveness (Spearman correlation: ρ < 0.22, p > 0.38). There were also qualitative differences among the sampled fruits. For instance, mango cultivars (*M. indica*) emit a diverse suite of terpenoid compounds (Fig 2A; Table S2), whereas guava (*P. guajava*) cultivars were enriched in short-chain aliphatics compounds, such as hexenol acetate and ethyl butyrate. Members of the *Prunus* group (plums, peaches, and nectarines) slightly differed in their scent composition, with plums and peaches emitting higher amounts of aliphatic alkanes (e.g., hexa- and heptadecane), while the nectarines emitted more sesquiterpenes and aromatic compounds (e.g., α-farnesene, and benzaldehyde, respectively). Banana (*M. acuminata*) scent was composed of aliphatic esters and short-chain compounds (e.g., isoamyl acetate, acetic acid), whereas plantain (*M. **×** paradisiaca*) scent was enriched in alkanes and monoterpene compounds (e.g., hexadecane and limonene). Both pear species emitted scents enriched in the sesquiterpene α-farnesene, but *P. communis* emitted more aliphatic esters, while *P. pyrifolia* emitted more sesquiterpenes (Fig 3; Table S2).

**Fig 2.**
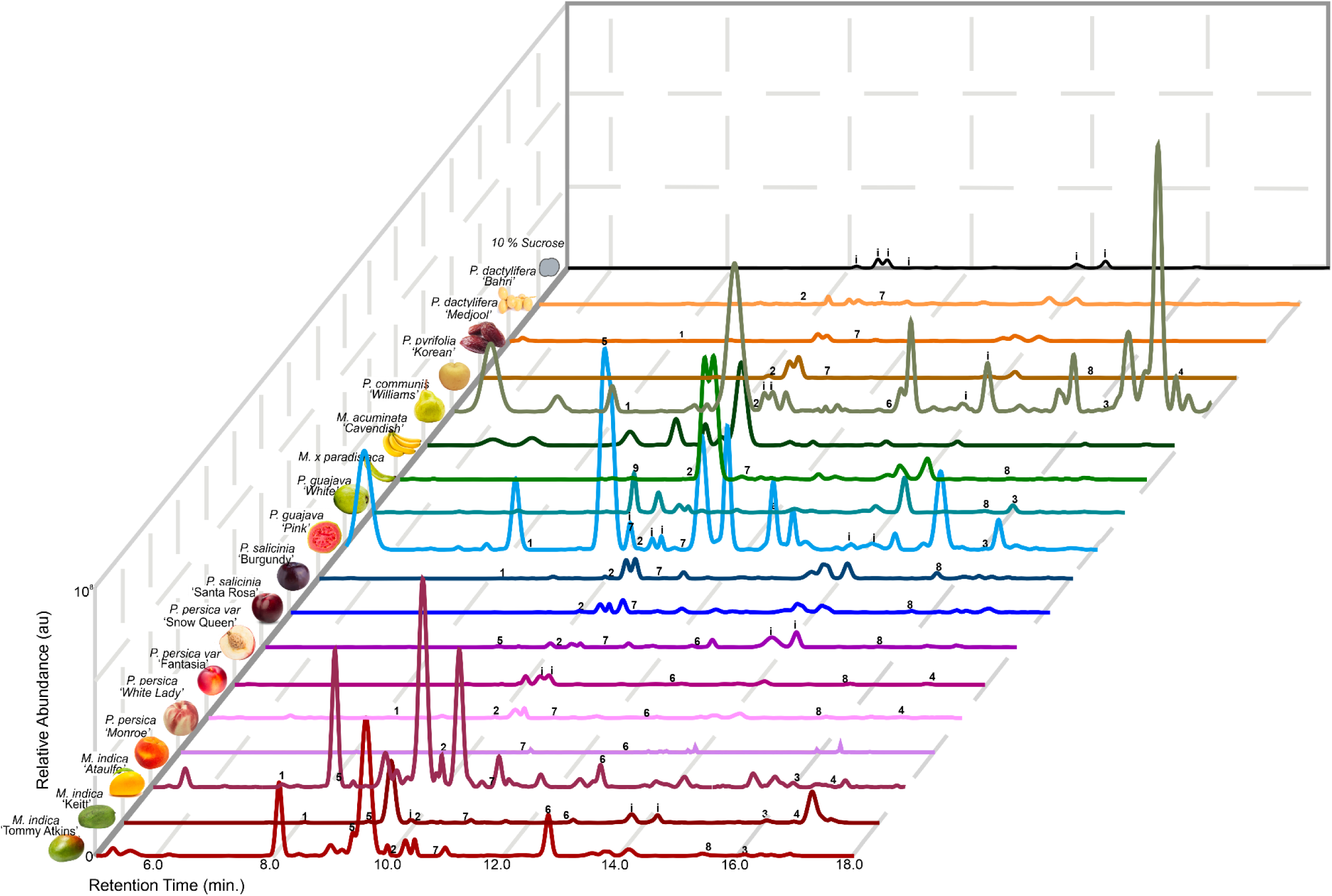
Ion chromatograms and chemical profiles of fruit scents. Representative Gas Chromatography-Mass spectrometry (GC-MS) ion chromatograms for each corresponding fruit and associated compounds of interest. Numbers indicate: 1) α-pinene, 2) Limonene, 3) Caryophyllene, 4) α-Farnesene, 5) Ethyl hexanoate, 6) Ethyl octanoate, 7) Nonanal, 8) Tetradecane 9) Hexen-1-ol-acetate (*E,Z*). Contaminants are denoted by *i*.

**Fig 3.**
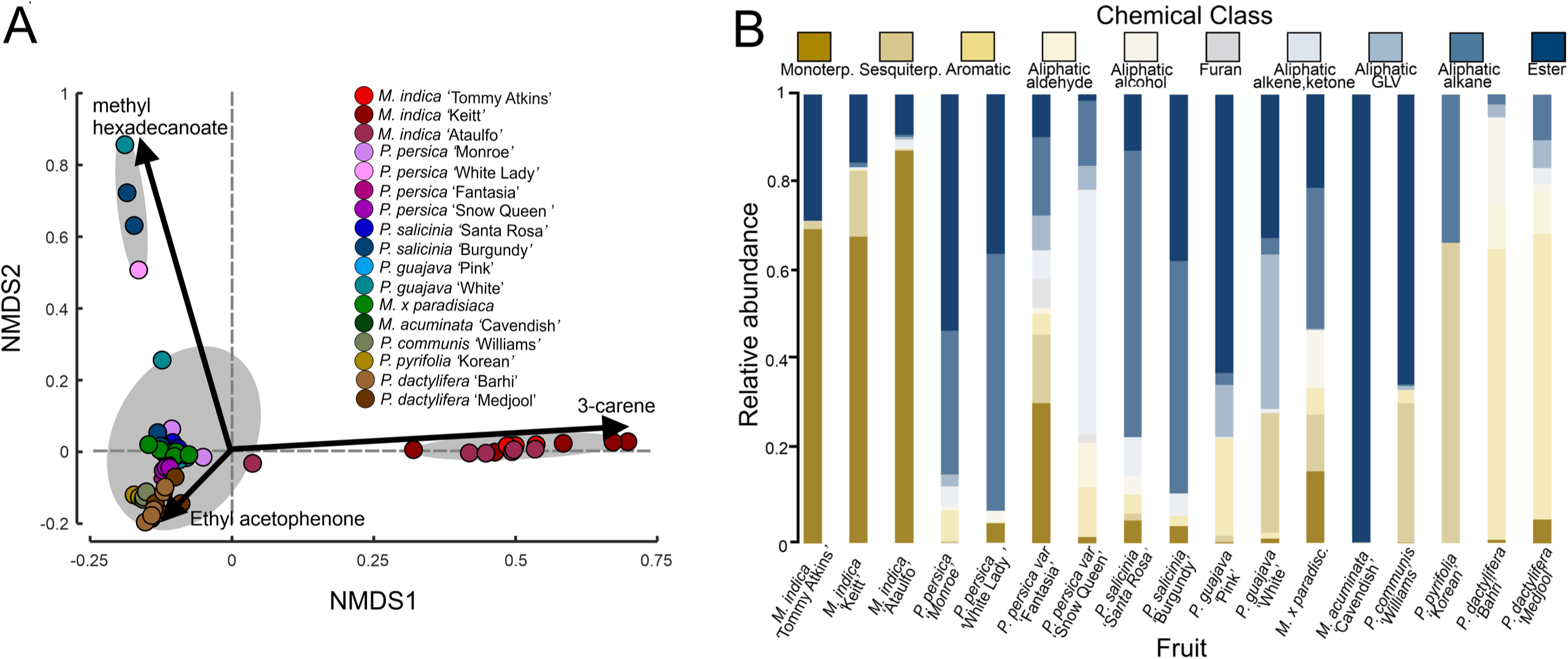
Chemical type proportions and multivariate analysis of fruit scents. **(A)** Nonmetric multidimensional scaling (NMDS) biplot of the chemical composition of all fruit groups and varieties are represented by color and corresponding shade. Analysis of the scent compositions are significantly different between fruits (ANOSIM: R = 0.8417, p = 0.001), with significantly different clusters denoted by gray ellipses. Labeled arrows denote the chemical compounds dominating the different axes. **(B)** Fruit scent profile demonstrating the ratio of different classes of compounds for each fruit group and variety, including: monoterpenes, sesquiterpenes, aromatics, aliphatic aldehydes, aliphatics alcohols, furans, aliphatic alkenes & ketones, aliphatic – GLVs (green leaf volatiles), aliphatic alkanes, and esters.

To analyze the variability generated by the 150 compounds across the 19 fruit species, varieties, and cultivars, we conducted a multivariate analysis (NMDS) using the proportion of compounds in the fruit scents (Fig 3). This analysis also found a significant difference between fruit scents (ANOSIM: R = 0.6963, p = 0.001). The attractive white nectarines (*Prunus persica nucipersica* ‘Snow White’) were close to the other *Prunus* species and varieties, and close to the other fruit species (Fig 3A). By contrast, the mango (*M. indica*) cultivars occupied a distinct area along NMDS1 and NMDS2. By plotting individual volatiles in the same NMDS space, we found that the monoterpene 3-carene was distributed along the NMDS1 axis, whereas the sesquiterpene, caryophyllene, the ester, isoamyl acetate, and the aliphatic compound, hexadecane, were distributed along the NMDS2 axis (Fig 3A). Between the closely related species and fruit cultivars, their proportions of compound types in the scents also showed significant variation, especially their proportion of monoterpenes. For instance, guava (*P. guajava*) cultivars differed in their terpenoid and aromatic compound proportions, nectarine (*P. persica nucipersica*) cultivars differed in their proportion of monoterpene and aromatic compounds, and mango (*M. indica*) cultivars differed in their relative amounts of terpenes (Fig 3B).

### Antennal olfactory responses to the fruit scents

The select preference of *Ae. aegypti* for the scents of certain fruits and fruit varieties motivated us to examine whether their antennae respond differently to those scents. We performed electroantennogram (EAG) recordings to measure the summed response of olfactory sensory neurons on the mosquito antennae to a panel of fruit scent extracts (Figs. 4, S1). Results from these experiments showed that fruit scent extracts evoked stronger EAG responses relative to the blank odor cartridge and solvent controls (Fig. 4; Kruskal-Wallis test: χ17,274 = 95.66, p < 0.0001). The mangos (*M. indica*), guavas (*P. guajava*), peaches (*P. persica*), nectarines (*Prunus persica var nucipersica*), and bananas (*Musa* spp.) evoked significantly stronger responses than the solvent control (Dunn-Sidak test: p < 0.02), but the plums (*P. salicinia*) and pears (*Pyrus* spp.) were not different from the controls (Dunn-Sidak test: p > 0.10). Among each pair of related species and cultivars, only the guava (*P. guajava*) cultivars evoked significantly different responses from each other, with the ‘White’ cultivar evoking stronger responses than the ‘Pink’ (Mann-Whitney U-test: *p* = 0.02). There was no significant correlation between the EAG responses and fruit scent emission rates (Spearman correlation: ρ = −0.48, p = 0.06), nor the EAG responses and the number of mosquitoes attracted to the scents (Spearman correlation: ρ = 0.05, p = 0.85).

**Fig 4.**
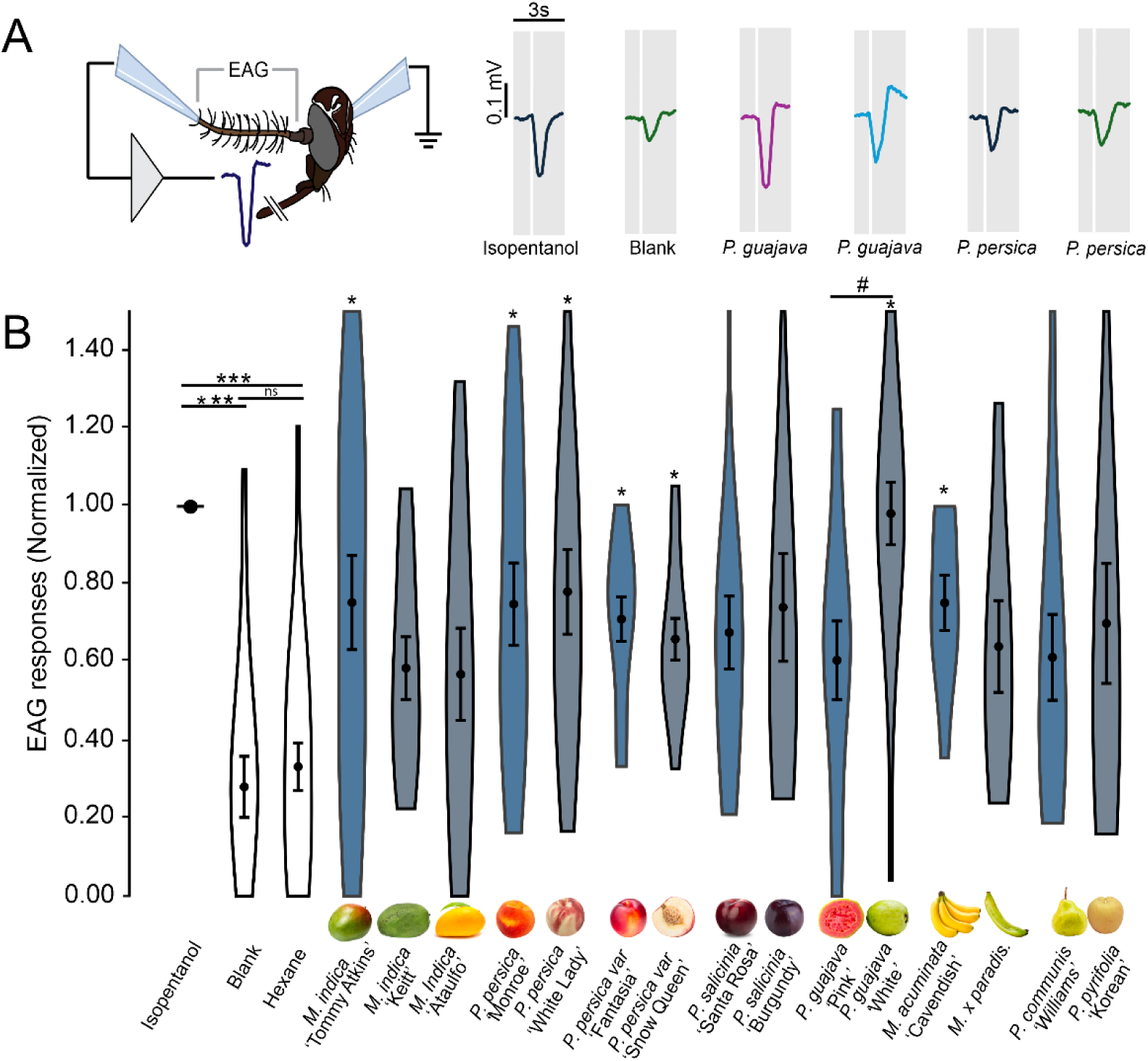
Electroantennogram responses to the fruit scents. (**A**) Experimental setup for the electroantennogram (EAG) experiments (image courtesy of M. Stensmyr). Traces are the individual EAG responses to an olfactory stimulation (0.5 sec.) showing the responses to isopentanol (+control), clean air blank (-control), guava varieties (*P. guajava*, ‘White’ and ‘Pink’), and nectarine varieties (*P. persica variety (var) nucipersica* ‘Fantasia,’ and ‘Snow Queen’). (**B**) Violin plots of EAG responses across the tested olfactory stimuli; the mean ± SEM for each stimulus is shown in each plot. Statistical analyses were performed with the normalized data (relative to isopentanol control in each preparation). Plots are the Mean ± SE (n = 8-25 mosquitoes/odor stimulus). Asterisks denote significant responses compared to the solvent control (Mann-Whitney U-test: *p* < 0.05). Comparing responses between varieties and closely related fruit species showed that only *P. guajava* varieties elicited significantly different responses (Mann-Whitney *U*-test: *p* = 0.001; denoted by #).

### Source of dominant volatiles in the fruit scent

To evaluate the potential sources of the different volatile organic compounds emitted from the fruits, we sampled the headspace of the peels (exocarp) and pulp (mesocarp) of three of the most attractive fruit species: red mango (*M. indica* ‘Tommy Atkins’), white nectarine (*P. persica nucipersica* ‘Snow Queen’), and banana (*M. acuminata* ‘Cavendish’)(Fig 5A). For all three fruits, there was a significant difference in the scent between the fruit exocarp and mesocarp (Kruskal-Wallis test: χ5,17 = 13.82, p = 0.01), with the exocarp emitting 3.7 to 8.9-fold higher levels of volatile organic compounds compared to the mesocarp (Fig 5B). These quantitative differences in scent emission were also reflected in differences in chemical composition and proportions between the parts of the fruit. Examples of these differences come from mango (*M. indica*), where the pulp lacked 3-carene, a dominant compound emitted from the peel and whole fruit, comprising up to 56% of the total scent emission (Figs 1B, 3B). The compound proportions also differed between the mango peel and pulp, with terpenes dominating the scent emissions of the peel and whole fruit, whereas ketones and short-chain alcohols were the dominant compounds in the scent of the pulp (e.g., cyclopentanone, 1-hexanol). These differences in the composition and compound proportions of the peel and pulp scents were also found in the nectarine (*P. persica nucipersica* ‘White’) and banana (*M. acuminata* ‘Cavendish’). The nectarine pulp scent was dominated by cyclopentanone, whereas the peel scent was dominated by fatty acid esters like ethyl hexanoate. Banana pulp scent included many different compounds, including 2-pentanol acetate and isopentyl isobutyrate, whereas its peel scent was dominated by esters like isoamyl butanoate and isoamyl acetate (Table S2).

**Fig 5.**
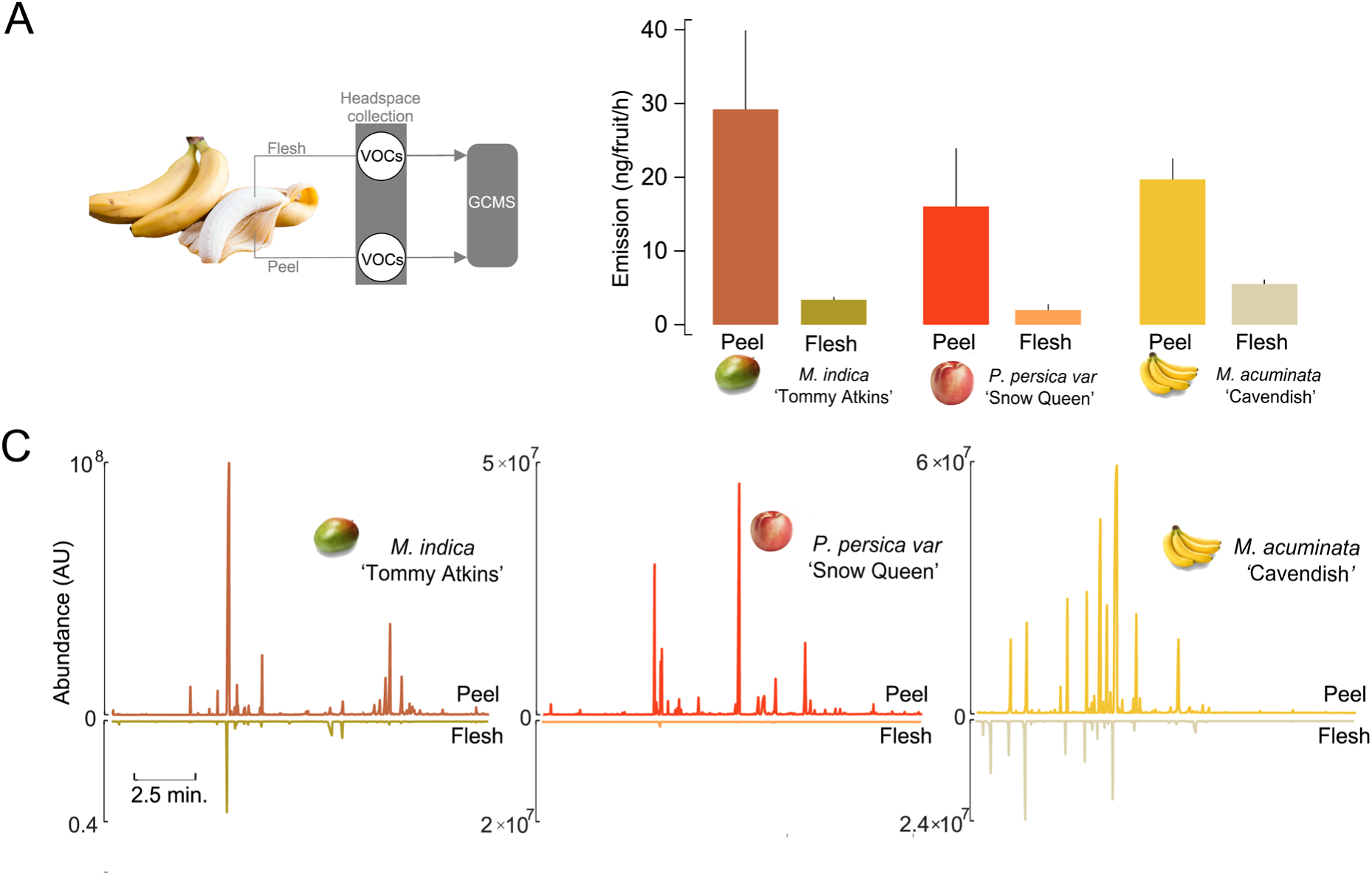
Volatile source emissions from fruits. (**A**) Schematic of the headspace collections from the exocarp (peel) and mesocarp (pulp). (**B**) Emission of peel and mesocarp of three fruits (*M. indica* ‘Tommy Atkins’*, P. perspica nucipersica ‘*Snow Queen’ (white nectarine), and *M. acuminata* ‘Cavendish’ (banana)). Bars are the mean ± SE. (**C**) Ion chromatograms of the peel and mesocarp of the three fruits: *M. indica* ‘Tommy Atkins’*, P. persica nucipersica ‘*Snow Queen’ (white nectarine) and *M. acuminata* ‘Cavendish’ (banana).

### Behavioral Effects of Altering the Compound Proportions in the Fruit Scents

The differences in the attractiveness of closely related fruits, and how those differences are reflected in the proportion of compounds in the scent, motivated us to ask if changing the proportion of a single compound was sufficient to decrease the attractiveness of the fruit scent to levels similar to the related variety, cultivar, or species. For both nectarines and mangos (*P. persica var. nucipersica*, and *M. indica*, respectively), the difference between an attractive versus non-attractive cultivar was reflected in the proportion of monoterpenes in the scent, particularly the compound ⍺-pinene. For both white nectarines and red mangos, we increased their ⍺-pinene emissions by using a glass vial filled with mineral oil and a specific concentration of ⍺-pinene, placing the vial immediately next to the fruit, allowing us to simulate the ⍺-pinene emissions of the yellow nectarine or yellow mango in the context of the other fruit cultivar’s scent (Fig 6). Similar to the behavioral experiments testing the attractiveness of the different fruit scents (Fig 1A), mosquitoes were exposed to traps with the fruits (with and without the ⍺-pinene) and control traps for 48 h.

**Fig 6.**
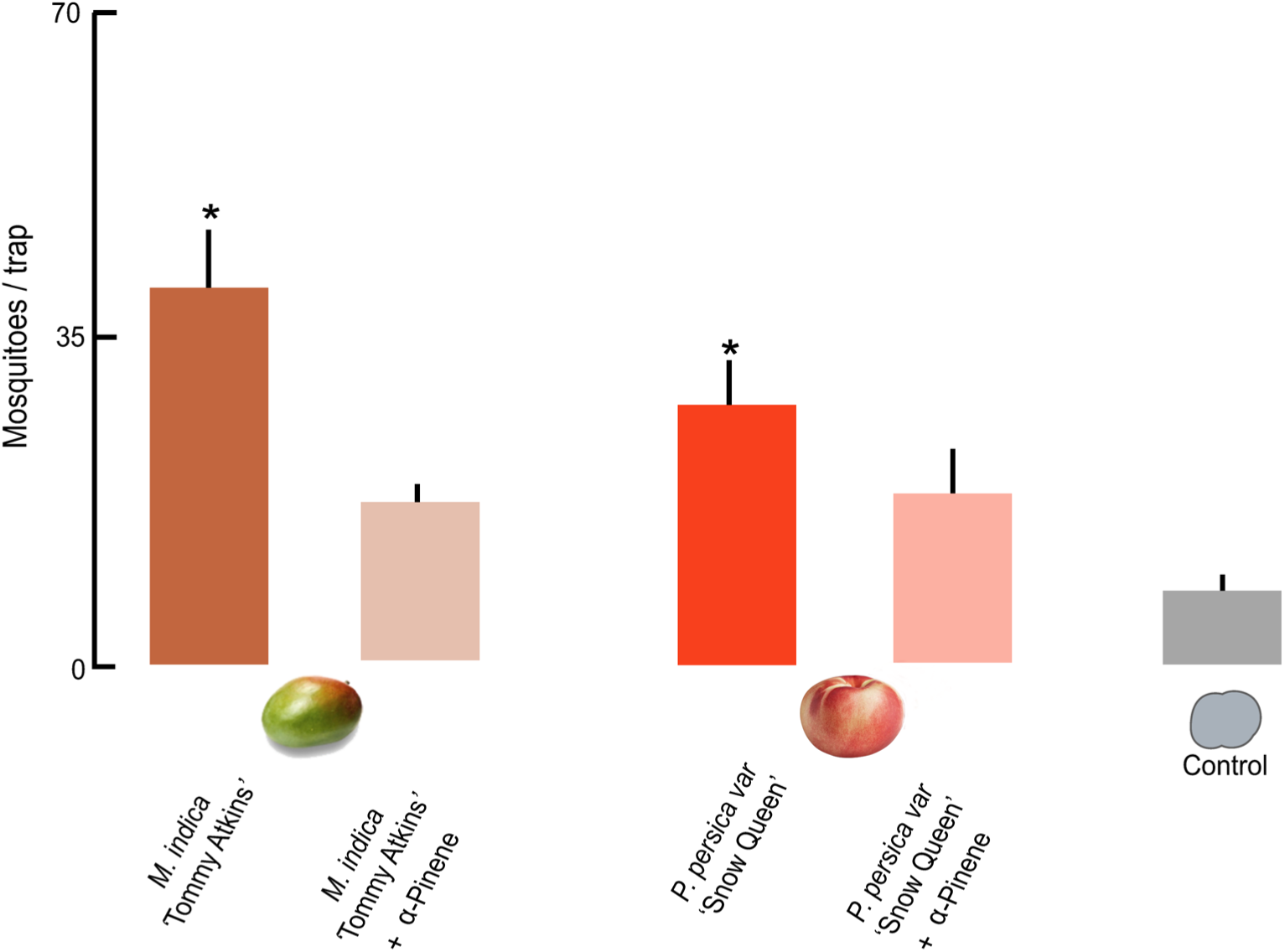
The effects of altering the compound proportions in *M. indica* and *P. persica nucipersica* scents. The number of mosquitoes attracted to traps emitting scents from *M. indica* (‘Tommy Atkins’) and *P. persica nucipersica* (‘Snow Queen’) fruits, with the addition of either a solvent control or α-pinene to simulate the proportion of α-pinene emitted from the less attractive varieties (*M. indica* ‘Ataulfo,’ and *P. persica nucipersica* ‘Fantasia’). In parallel, within-cage controls (10% sucrose) were run. Bars are the mean ± SE. Asterisks denote a significant difference between the fruit treatments (Mann-Whitney U-test; *=*p*<0.05).

For both red mango and white nectarine, results showed that fruits with increased ⍺- pinene emissions were significantly less attractive than the solvent (control) fruits (Mann-Whitney *U*-tests: *p* < 0.01). Moreover, comparing the level of attractiveness of the red mango scent spiked with ⍺-pinene to the yellow mango cultivar (*M. indica* ‘Ataulfo’), which has higher levels of ⍺-pinene emissions (Fig. 1B), showed no significant difference (Mann-Whitney U-test: *p* = 0.07). A similar result was found comparing the white nectarine spiked with ⍺-pinene to the yellow nectarine, which naturally emits higher levels of ⍺-pinene; there were no significant differences in their level of attraction (Mann-Whitney *U*-test: *p* = 0.43). Across all treatments, there was no difference between the number of mosquitoes in the control traps (Mann-Whitney U-tests: *p* > 0.27), suggesting that the low emissions of ⍺-pinene did not affect the locomotion or flight responses of the mosquitoes. Taken together, these results suggest that a change in the proportion of a single compound in the fruit scent, simulating the chemical composition of the less attractive cultivar, can significantly reduce the attractiveness of a fruit.

## Discussion

Motivated by the dearth of studies examining how differences in the scents of plant sugar sources influence mosquito attraction, we examined the preference of *Ae. aegypti* mosquitoes to the scents of closely related fruits and their cultivars. Our results show that both male and female *Ae. aegypti* mosquitoes have attractive preferences for specific fruit species and varieties, including mangos (*M. indica*), nectarines (*P. perspica var. nucipersica*), guavas (*P. guajava*), and bananas (*M. acuminata*), whereas other fruits (*P. communis, C. limon*) were not attractive. Although the chemical profile can differ between disparate species—for example, *Musa spp.* compared to *Mangifera*—the differences within species predominantly reflect changes in the proportion of compounds in the scent. Similar to prior work testing different flower scents (Lahondère et al., 2020), our results show that the proportion of compounds in the fruit scents can have strong behavioral effects.

Mosquitoes are attracted to diverse flowers, fruits, and honeydew as plant sugar sources (Athen et al., 2020; Lahondère et al., 2020; Peach et al., 2019; Yalla et al., 2023). However, various studies, including those in the laboratory, semi-field, and field, have shown that plant sugar sources can be differentially attractive to mosquitoes, with some plants highly attractive and others eliciting little to no attraction (Gary Jr and Foster, 2004; Müller et al., 2011). In an elegant series of experiments, Nyasembe et al. (2018) examined the putative feeding preferences of field-caught mosquitoes in Kenya using DNA bar-coding and found that *Anopheles gambiae s.s.* mosquitoes may predominantly have fed on a subset of plants in the environment, such as *Senna alata* (Fabaceae), *Ricinus communis* (Euphorbiaceae), and *Parthenium hysterophorus* (Asteraceae), whereas *Ae. aegypti* mosquitoes may have fed on *Senna uniflora* (Fabaceae) and *Hibiscus heterophyllus* (Malvaceae)(Nyasembe et al., 2018). Similar results have been shown in different mosquito species may be feeding from diverse flowering plants (Lahondère et al., 2020; Manda et al., 2007; Müller et al., 2010; Müller et al., 2011; Yu et al., 2017). Research examining the preferences of *Culex pipiens pallens* mosquitoes indicated that they are differentially attracted to several flowering plant species, including *Tagetes erecta* (Asteraceae) and *Catharanthus roseus* (Apocynaceae)(Yu et al., 2017). *Aedes albopictus* adults were differentially attracted to *Tamarix chinensis* (Tamaricaceae), *Ziziphus spina-christi* (Rhamnaceae), *Prosopis farcta* (Mimosaceae), and other plant families (Müller et al., 2011). The diversity in plant species and families used as sugar sources makes it difficult to identify specific plant groups mosquitoes may feed on, and instead may reflect similarities in the chemical profiles of the flowering plants, or local plant abundances that mosquitoes can adaptively utilize.

In contrast to the growing body of work using flowering plants, research on mosquito preference for intact fruit scents has received comparatively less attention. Examples include the mosquito *Culex pipiens pallens*, which showed attraction to the scents of peach and melon (*Amygdalus persica* and *Cucumis melo*, respectively) but was less attracted to pear (*Pyrus bretschneideri*)(Yu et al., 2017). In field trials, male and female *Aedes albopictus* mosquitoes showed attraction to sabra and figs (*Opuntia ficus indica* and *Ficus carica*, respectively) but not undamaged pomegranate (*Punica granatum*)(Müller et al., 2011). Research from this current study shows that attraction can also vary between closely related fruit species and variety, and although we only tested male and female mosquitoes of one species (*Ae. aegypti*), previous work in other mosquito species, such as *An. gambiae* has shown similar attraction to some of the tested species but different cultivars, including *M. indica* ‘Kent’ (Meza et al., 2020).

This strong preference by mosquitoes for specific fruit scents also raises the question about the relatedness and differences in the scent profile between plant sugar sources. Although the chemical composition of the sources of plant sugar can differ, many of the attractive scents share the presence of compound types in their profile, including various isomers of pinene, myrcene, terpinolene, linalool and linalool oxide, and caryophyllene (Lahondère et al., 2020; Nikbakhtzadeh et al., 2014; Tenywa et al., 2017). Other compound types, including aliphatic aldehydes and esters, have also been shown to be important for mosquito detection of plant sugar sources (McGovern et al., 1970). Similar compounds are found in the headspace of many fruits tested in this study, including mango, peach, and nectarine (Figs. 2 and 3). The banana (*M. acuminata*) is another fruit that is attractive to *Ae. aegypti* mosquitoes and emitted a scent that was dominated by aliphatic ester compounds, including 2-pentyl acetate, 3-methylbutyl acetate, and 3-methylbutyl butanoate (Table S2), some of which were also emitted by the attractive mango and guava fruits. Nonetheless, across these similarities, the differences in scent compositions between closely related species and varieties may provide insight into the compounds that decrease the attractiveness of the fruit scents. For example, red mango and white nectarine (*M. indica* ‘Tommy Atkins,’ and *P. persica nucipersica* ‘Snow Queen’, respectively) emitted lower amounts of monoterpenes, including α-pinene, compared to the varieties and species that were less attractive. Increasing the concentration of α-pinene in the scents decreased their attractiveness (Fig 6). Beyond monoterpenes, the black plum (*P. salicinia* ‘Burgundy’) emits higher levels of esters and aliphatic aldehydes, known attractants to mosquitoes (Bosch et al., 2000; Takken and Knols, 1999), compared to the red plum (*P. salicinia* ‘Santa Rosa’), whereas the yellow peach emits higher levels of benzenoid compounds, such as the benzaldehyde, acetophenone, and 2-ethyl benzoate, than the white peach (*P. persica* ‘White Lady’). These benzenoid compounds have been implicated as both attractants (acetophenone)(Afify and Potter, 2020) and repellents (benzaldehyde)(Zhang et al., 2022) in mosquitoes, and the concentration and proportion of these compounds in the scent may be critical for the valence of the mosquito’s behavior. Besides the specific compounds in the scents, the differences in attraction between cultivars and varieties may be related to their scent profiles and the proportion of compounds in the scents (Figs. 2 and 3). An important aspect of this study is the Porapak Q adsorbent used to collect fruit scents. This adsorbent, although ideal for diverse volatile types, will not collect small molecular weight compounds (<100 Da), nor efficiently collect polar compounds such as fatty acids. Future work will need to use alternate adsorbent methods, such as solid-phase microextraction fibers with polydimethylsiloxane/divinylbenzene coating, to allow the capture and identification of polar and semi-volatile compounds in the scents, as well as characterize the different chiral compounds in these scents. Despite these potential caveats, our work quantified a diverse panel of compounds in the fruit scents and showed that manipulating the concentration of a single compound in an attractive headspace was sufficient to lower the fruit’s attractiveness to levels similar to its non-attractive cultivar.

The differences in behavioral attraction were not reflected in the antennal olfactory responses to the fruit scents. Results from our EAG experiments showed that the mosquito antennae evoked strong responses to many of the fruit scents compared to the negative control, but that only one pair of varieties, from the guava (*P. guajava*), elicited significantly different responses (Fig. 4). Similar EAG responses with differing behavior may reflect the downstream processing in the mosquito’s brain. Prior work has shown that neuropil in the mosquito antennal lobe are sensitive to subtle differences in the composition and proportion of compounds in closely related floral scents (Lahondère et al., 2020) – similar results could be occurring in the neural coding of the fruit scents. An important missing gap from our current work is the lack of identifying the bioactive compounds within the complex fruit scents. Previous work using Gas Chromatography with Electroantennogram Detection (GC-EAD) has shown that mosquitoes are responsive to a variety of compounds in the scents of plant sugar sources. For example, Nyasembe et al (2018) found that *Ae. aegypti*, *An. gambaie*, and *Aedes mcintoshi* mosquitoes detected a similar set of monoterpenes (linalool, linalool oxide, *β*-myrcene, and *β*-ocimene) in the floral scents. Qualitatively analogous results were found by Lahondere et al. (2019), where *Ae. aegypti, Anopheles stephensi* and *Aedes communis* mosquitoes responded to similar monoterpene and aliphatic aldehyde compounds, such as linalool, lilac aldehyde, *β*-myrcene, *β*-ocimene, nonanal, and decanal. Future research will be needed to identify the fruit volatiles that are detected by the mosquitoes (via Gas Chromatography coupled Electroantennogram Detection, or GC-Single Sensillum Recording) to determine if the same compounds identified here in this study are detected across different mosquito species, and how the scents are encoded in the brain.

Attractants incorporating fruit scents to attract mosquitoes are thought to be an important control intervention when used with existing approaches like bed nets and insecticides (Njoroge et al., 2023). In limited field trials in Mali, Africa, lures based on Attractive Toxic Sugar Baits (ATSBs^TM^) were able to decrease the number of mosquitoes bearing malaria pathogens (Traore et al., 2020). However, more recent trials using ATSBs^TM^ have shown limited efficacy (Wagman et al., 2024), which raises the question of what might be causing these changes. Mosquito lures like ATSBs^TM^ often use fruit syrups or fermented fruit juices combined with insecticides to attract and kill feeding mosquitoes (Torto and Tchouassi, 2024; Traore et al., 2020), and variations in the sources of these syrups, such as using different cultivars, could potentially affect their attractiveness and efficacy (Fig 1B). Syrups and juices also do not incorporate the peel, the dominant source of volatiles (Thiruchelvam et al., 2020). In the natural setting, mosquitoes will be attracted to the compounds emitted from the fruit and damaged fruit on the ground, all providing a source of higher concentrations of scent compounds and occurring at their natural proportions (Joseph, 1970). Formulating peel-derived lures or using artificial odors that mimic attractive sugar sources could increase the attractiveness and longevity of the traps while decreasing sources of variation in the lure’s attractiveness.

Beyond the compounds emitted from the peel, and beyond the mosquito’s sense of smell, plant sugar sources provide other sensory cues that may attract mosquitoes. For example, the fruits emit high levels of water vapor that attract foraging mosquitoes (Grierson and Wardowski, 1978; Laursen et al., 2023), as well as providing a visually contrasting and spectrally-rich display of the sugar source. Once contacting and tasting the fruit, gustatory stimuli such as sugars, phenolics, terpenes, and other antioxidant compounds (Saini et al., 2022), could be potentially detected by the mosquito (Baik and Carlson, 2020). The relative contribution of these other sensory cues in mediating attraction and feeding on plant sugar sources remains untested. By contrast, a growing number of studies have shown the importance of multiple sensory cues in mediating attraction to blood hosts (McMeniman et al., 2014), including the combination of CO_2_, heat, skin odor, water vapor, and/or visual displays (Alonso San Alberto et al., 2022; Chandel et al., 2024; Giraldo et al., 2023; Laursen et al., 2023). Future work will be needed to examine these in more detail for sugar sources, and how the mosquito nervous system detects and processes complex olfactory and multimodal information.

## Acknowledgments

We are grateful for the discussions and advice provided by A. Rouyar, C. Ruiz, S. Jandu, I. Vieira Coutinho Abreu Gomes, O. Akbari, and J. Pitts. We thank B. Nguyen for mosquito rearing and care, and L. Schwyhart for assistance in behavioral experiments.

## Author contributions

Conceptualization: MLN and JAR; Methodology: MLN, GMT, and JAR; Formal analysis: MLN and JAR; Investigation: MLN, GMT, and JAR; Data curation: MLN and JAR; Writing - review & editing: JAR, MLN, and GMT; Visualization: MLN and JAR; Supervision: JAR; Project administration: JAR; Funding acquisition: JAR.

## Data availability

Data are available from Mendeley Data; code is available at https://github.com/riffelllab.

## Supplementary Information

**Supplementary Information Table S1.** Fruit details and origins.

**Supplementary Information Table S2.** GCMS analyses of fruit scent composition and emission rates.

**Figure S1. Electroantennogram responses (µV) across the tested olfactory stimuli.** Boxes are the Mean ± SE (n = 8-25 mosquitoes/odor stimulus).

